# Data-informed model reduction for inference and prediction from non-identifiable models

**DOI:** 10.1101/2025.03.20.644462

**Authors:** Matthew J. Simpson

## Abstract

Many mathematical models in the field of theoretical biology involve challenges relating to parameter identifiability. Non-identifiability implies that different combinations of parameter values lead to indistinguishable solutions of the mathematical model. This means that it is difficult, and sometimes impossible, to explain the mechanistic origin of observations using a non-identifiable mathematical model. A standard approach to deal with structurally non-identifiable models is to use reparameterisation, which typically focuses on the structure of the mathematical model without accounting for the impact of noisy, finite data. We present and explore a simple computational approach for model reduction, via likelihood reparameterisation, that can be applied to both structurally non-identifiable and practically non-identifiable problems. We construct simplified, identifiable mathematical models that enable model-based predictions for a range of continuum models based on different classes of commonly-used differential equations. Through a series of computational experiments, we illustrate how to deal with a range of noise models that relate the solution of the mathematical model with noisy observations. A key focus is to illustrate how computationally efficient model-based predictions can be made from reduced models.

## 1 Introduction

Parameter identifiability is a major challenge in the field of theoretical biology [1], and is often considered in terms of *structural identifiability* and *practical identifiability*. In this work we will deal with both forms. Structural identifiability can be thought of as a property of a mathematical model, and refers to whether model parameters are uniquely determined in the hypothetical situation where we have access to an infinite amount of ideal noise-free data [2, 3, 4, 5, 6, 7]. Practical identifiability, in contrast, is a joint property of a particular mathematical model with a particular dataset, and refers to the extent to which model parameters can be estimated in the face of finite, noisy, imperfect data [5, 8, 9, 10, 11, 12, 13, 14]. An obvious way of addressing practical non-identifiability is to improve the quality and/or quantity of the dataset so that parameters can be estimated with increased precision [15, 16]. Classical approaches for dealing with structural non-identifiability involve model reduction by seeking a reparameterisation that may reduce the number of parameters [17, 18, 19, 20, 21, 22, 23]. Throughout this work we will also refer to the mathematical model as a *process model*, since the mathematical model is an abstract representation of the biological process of interest.

Parameter identifiability is a critical challenge for the theoretical biology community partly because model development often takes place without quantitatively comparing model predictions with experimental data. The lag between theoretical model development and practical model application means that challenges of parameter identifiability are sometimes discovered *post hoc* as long-established models are subsequently calibrated using new datasets [15, 16].

In this work we outline a computationally-based, data-informed workflow for model reparameterisation. As we will demonstrate, these methods can be applied to both structurally and practically non-identifiable problems. In essence, the approach involves finding a computationally-derived reparameterisation of the original model to give a reduced model. The reduced model involves less parameters than the original model, and the parameters in the reduced model are identifiable.

Following the approaches used in the sloppy analysis of mathematical models [24, 25, 26, 27, 28, 29], we reparameterise the model using an eigendecomposition of the observed Fisher Information evaluated at the maximum likelihood estimate. This gives rise to a transformation from the *original* model parameters into a new set of *eigenparameters* which facilitates model reduction and simplification. The decomposition provides a list of eigenparameters ordered in terms of their identifiability [30]. For the tutorial style examples that we present, the observed Fisher Information for structurally non-identifiable models is rank deficient, whereas for practically non-identifiable models the observed Fisher Information has full rank. Regardless of the form of non-identifiability, the eigendecomposition provides a means of combining parameters in a reduced identifiable mathematical model. For the reduced model we explore both estimation and prediction, illustrating how reduced models can be used to explore the extent to which profile likelihood-based confidence sets for model parameters can be combined efficiently to give an overall prediction confidence set that, as we show, can provide an accurate approximation to the exact prediction confidence set obtained by sampling the full likelihood.

A key aim is to illustrate how these concepts of data-informed reparameterisation and model reduction can be applied to a range of continuum mathematical models with different forms of observational data. In particular, we provide illustrations that involve modelling with ordinary differential equations (ODE), taking the form of both initial value problems (IVP) and boundary value problems (BVP), as well as working with mathematical models in the form of partial differential equations (PDE). In particular, our computational examples illustrate how the same framework can be used to deal with a range of noise models that relate the solution of the mathematical model with noisy observations (e.g. additive Gaussian noise, multiplicative log-normal noise, multinomial models for count data). Software to replicate our results are provided in the formal of Jupyter notebooks that are written in the open access Julia language.

## 2 Methodology

### 2.1 Likelihood-based estimation and identifiability

We employ a likelihood-based approach for estimation and identifiability analysis, and the details of the likelihood function will differ depending on the form of the process model we work with, including ODE, BVP and PDE-based mathematical models. The evaluation of the likelihood function also depends upon details of the noise model that relates individual measurements to the output of the process model, and in this work we will deal with a range of noise models including additive Gaussian, multiplicative log-normal, and multinomial noise models that are useful for working with count data in population biology. Regardless of these specific details, we always work with a loglikelihood function that we write as 𝓁(***θ*** | ***y***^obs^), where ***θ*** is a vector of length *K* containing the model parameters, and ***y***^obs^ is a vector of observed data. The form of the data will depend upon the details of the process model and experimental design, and we will explain these differences in each example covered.

Applying maximum likelihood estimation (MLE) gives the best-fit set of parameters [31, 32],

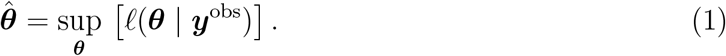

All numerical optimisation computations in this work use the Nelder-Mead algorithm with simple bound constraints [33], and all computations are performed using default stopping criteria within the well-known NLopt family of numerical optimisation routines [33]. Given the MLE estimate, we then work with the normalised loglikelihood function 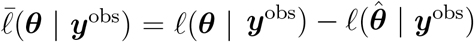, so that 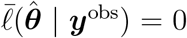. Throughout this work we consider 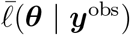 to be a function of ***θ*** for a fixed dataset. Once we have established the normalised loglikelihood function we are able to define asymptotic confidence sets by identifying a likelihood-based threshold, 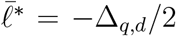, where Δ_*q,n*_ refers to the *q*th quantile of the *χ*^2^ distribution with *n* degrees of freedom [34, 35]. This threshold allows us to define asymptotic confidence sets defined implicitly by the values of ***θ*** satisfying 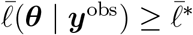.

To assess parameter identifiability we take a profile likelihood-based approach [36, 37] by partitioning ***θ*** into interest parameters *ψ*, and nuisance parameters *ω*, so that ***θ*** = (*ψ, ω*)^⊤^. For all calculations we either take the interest parameter to be a single parameter which allows us to construct various univariate profile likelihood functions, or a pair of parameters which allows us to construct various bivariate profile likelihood functions [37]. Regardless, for a set of data ***y***^obs^, the profile loglikelihood for the interest parameter *ψ* given the partition (*ψ, ω*)^⊤^ is

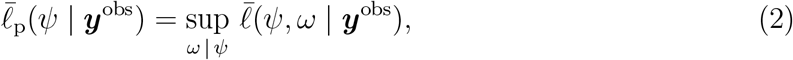

which implicitly defines a function *ω*\* (*ψ*) of optimal values of *ω* for each value of *ψ*, and defines a surface with points (*ψ, ω*\*(*ψ*)) in parameter space. For univariate profiles, 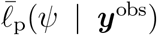 is a function of one parameter which can be plotted and visualised in the usual way. For bivariate profiles, 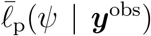 is a function of two parameters that can be visualised as a contour plot, heat map or similar. As for calculating the MLE, profile likelihoods are computed using the Nelder-Mead algorithm to evaluate the necessary numerical optimisation problems [33], and in all cases the iterative optimisation calculations can be started using the appropriate components of 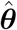. Regardless of whether we consider a univariate or bivariate profile likelihood function we evaluate 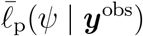 on a uniform discretisation of the interest parameter, *ψ*.

### 2.2 Data-informed reparameterisation

Inspired by the literature on model sloppiness [24, 25, 26, 27, 29], we consider the observed Fisher Information at the MLE,

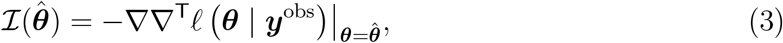

which is a *K* × *K* square matrix whose elements we compute using automatic differentiation [38, 39]. We compute the eigenvalues and eigenvectors, *λ*_*k*_ and **v**_*k*_ for *k* = 1, 2, 3, …, *K*, respectively of this square matrix using standard linear algebra routines in Julia. Since the eigenvectors are mutually orthogonal, the eigenparameters can be written as linear combination of the original parameters

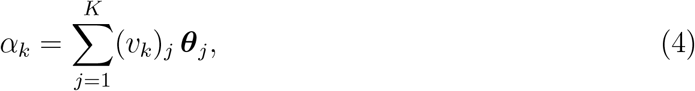

where (*v*_*k*_)_*j*_ is the *j*th element of the *k*th eigenvector **v**_*k*_, meaning each eigenparameter is simply a linear combination of the original model parameters. In all our results we work with normalised eigenvectors so that (*v*_*k*_)_*j*_ ∈ [−1, 1]. As we will demonstrate, for structurally non-identifiable models 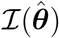 is rank deficient. Under these conditions, the eigenvectors associated with zero eigenvalues span the non-identifiable parameter space, and the eigenvectors associated with non-zero eigenvalues span the identifiable parameter space [17, 18, 40]. For structurally non-identifiable models, the eigendecomposition will lead to a reduced model where the number of eigenparameters is less than the number of original parameters.

For certain problems, we will work in a log parameterisation of the loglikelihood function and proceed in precisely the same way, leading to eigenparameters that take the form of a linear combination

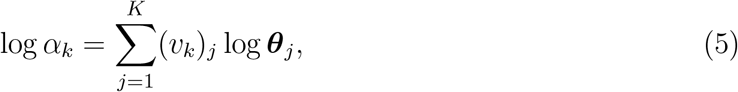

which can be re-written so that the eigenparameters are expressed as a product or quotient of original parameters raised to a power given by the elements of the relevant eigenvector. As before, for structurally non-identifiable models 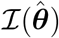 is rank deficient and this reparameterisation leads to a reduced model where the number of eigenparameters is less than the number of original parameters. Again, the eigenvectors associated with zero eigenvalues span the non-identifiable parameter space, and the eigenvectors associated with non-zero eigenvalues span the identifiable parameter space [17, 18, 40].

This approach to reparameterisation can be applied either directly or approximately, and we will illustrate both approaches. The direct application of this approach involves applying the relationships between ***α*** and ***θ***, as stated above. It is also possible to apply these relationships approximately when the elements of **v**_*k*_ are sufficiently close to zero. As we will demonstrate, in these cases terms involving these smaller elements can be omitted from the summation [27]. This approximation can lead to a reduced model that involves a combination of original and eigenparameters and, as we will demonstrate, this can be useful in the commonly-encountered situation where some parameters are well-identified by the available data whereas other parameters are poorly identified.

### 2.3 Likelihood-based prediction intervals

Likelihood-based estimation methods, described above, allow us to quantitatively determine how noisy data observations lead to both best-fit parameter estimates as well as asymptotic confidence sets of parameters [34]. We can determine how variability in ***y***^obs^ propagates into variability in ***θ***. With this information we then determine prediction intervals [41, 42] which provides a framework to understand how variability in ***θ*** impacts variability in the prediction of the mathematical model of interest. Throughout this work we will consider two types of prediction intervals [43, 44]: (i) exact prediction intervals constructed using the full loglikelihood function function 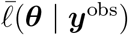; and, (ii) approximate prediction intervals constructed using *K* univariate profile loglikelihood functions, 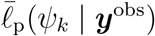, where *ψ*_*k*_ is the *k*th scalar interest parameter, and *k* = 1, 2, 3, …, *K*.

Exact prediction intervals are constructed using rejection sampling to draw parameter samples from the full loglikelihood function. Using this approach we obtain *M* samples, ***θ***_*m*_ for *m* = 1, 2, 3, …, *M* that lie within the 95% confidence region where 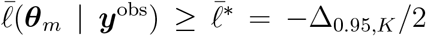 [34, 35]. For each of the *M* parameter samples, we solve the process model of interest to give a single model prediction (e.g. *x*(*t*)), which can be interpreted as the *mean trajectory* according to the relevant noise model [37, 43]. The width of the distribution underlying the noise model can be incorporated by computing the 5% and 95% quantiles of the associated noise model, which we denote *x*_0.05_(*t*) and *x*_0.95_(*t*), respectively. With this information we compute a single prediction interval, *x*(*t*) ∈ [*x*_0.05_(*t*), *x*_0.95_(*t*)]. Repeating this calculation for each ***θ***_*m*_, and taking the union of the resulting prediction intervals over the *M* parameter values gives us the exact prediction interval that we report in this work. This process requires *M* to be sufficiently large, and the choice of *M* depends upon the dimensionality and shape of 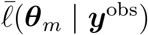. For simplicity we use *M* = 1000 throughout, noting that additional computation explorations (not shown) illustrate that further increasing *M* leads to visually-indistinguishable prediction intervals for the problems we consider.

For pedagogical reasons, in this work we restrict ourselves to relatively straightforward problems with *K* = 3 or *K* = 4 parameters only. Working with a small number of unknown parameters allows to make two important points. First, we show that issues of parameter identifiability are often encountered with very simple mathematical models, and that identifiability issues are not necessarily limited to overly complicated mathematical models. Second, since we are dealing with a set of computationally tractable problems, we are able to construct exact prediction intervals by using brute force rejection sampling on the full loglikelihood function. For more computationally expensive problems with a larger number of unknown, potentially correlated parameters, it can be computationally infeasible to sample from high dimensional likelihood functions. This motivates us to work with approximate profile-wise prediction intervals that can be constructed with a reduced computational cost.

Approximate prediction intervals are constructed using univariate profile loglikelihood functions, 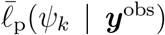, where *ψ*_*k*_ is the *k*th scalar interest parameter, for *k* = 1, 2, 3, …, *K*. Using rejection sampling we obtain *M* samples of *ψ*_*k,m*_ and *ω*_*k,m*_ for which 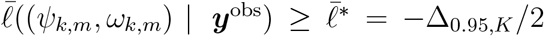 for all *k* = 1, 2, 3, …, *K*. As for the exact method (described above), for each of the *M* parameter combinations we solve the relevant process model, and use the 5% and 95% quantiles of the relevant noise model to compute a prediction interval, *x*(*t*) ∈ [*x*_0.05_(*t*), *x*_0.95_(*t*)]. We repeat this process for each of the *M* values of ***θ***_*m*_, and again for the remaining *K* − 1 univariate profile likelihood functions. Taking the union over the *M* × *K* prediction intervals gives the approximate prediction interval that we report. The re-parameterised loglikelihood functions lack any correlation structure which means that this approach can be accurate for modest *M*. Here we work with *M* = 50 [37].

## 3 Results and Discussion

We consider three examples that comprise different combinations of continuum mathematical models and different noise models relevant across a range of problems in mathematical biology. These include: (i) a structurally non-identifiable ODE-model with additive Gaussian noise; (ii) a structurally non-identifiable BVP with log-normal noise; and, (iii) a practically non-identifiable coupled PDE model describing experimentally observed count data using a multinomial noise model.

### 3.1 Production-decay model

To provide an illustration of our approach we consider a simple population dynamics model, where individuals within the population evolve according to a production-decay process,

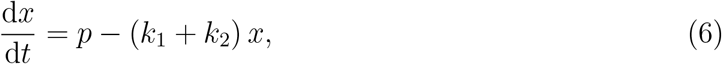

where *x*(*t*) ≥ 0 is the population density (e.g. population of molecules or a population of biological cells), *p* > 0 is a constant rate of production, and *k*_1_ > 0 and *k*_2_ > 0 are rates of decay. This simple model, with exact solution *x*(*t*) = *x*_s_ + (*x*(0) − *x*_s_) exp (−*t* [*k*_1_ + *k*_2_]), where 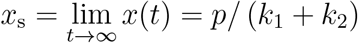, can be interpreted in many ways. For example, if *x*(*t*) represents a density of biological cells, the dynamics of the population is driven by cells being produced at a constant rate *p*, and cells undergo two different forms of linear decay at rates *k*_1_ and *k*_2_, respectively. In the context of modelling a population of cells, *k*_1_ could represent a rate of cell death due to apoptosis, and *k*_2_ could represent a rate of cell death due to necrosis. There are many other biologically relevant interpretations of this simple but insightful mathematical model.

The inference problem we consider is to estimate ***θ*** = (*k*_1_, *k*_2_, *p*)^⊤^ from a particular set of finite noisy data, ***y***^obs^(*t*). Before proceeding, we highlight that it is clear from the form of Equation (6), and its solution, that the mathematical model is structurally non-identifiable as the dynamics of the solution are controlled by the *sum* of the decay rates *k*_1_ + *k*_2_ and the production rate *p*. Since there are infinitely many choices of ***θ*** = (*k*_1_, *k*_2_, *p*)^⊤^ that give rise to the same values of (*k*_1_ + *k*_2_, *p*), the mathematical model is structurally non-identifiable. The same result can be obtained using structural identifiability software, such as DAISY [2], GenSSI [6] or STRIKE-GOLDD [4], in which systems of first-order ODE models are analysed in terms of Lie derivatives to understand the input-output relationship.

To explore the consequences of this we consider data generated with fixed initial conditions *x*(0) = 100, and true parameter values ***θ*** = (*k*_1_, *k*_2_, *p*)^⊤^ = (0.1, 0.1, 1.0)^⊤^. With this solution we construct ***y***^obs^ by recording 11 equally-spaced observations of *x*(*t*) across the interval 0 ≤ *t* ≤ 20, where each measurement is corrupted with additive Gaussian noise with *σ* = 3, which can be written as

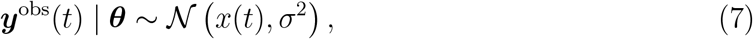

so that the mean of the noise model is given by the solution of the process model, and the variability in measurements is described by a constant variance, *σ*^2^, which we treat as a known constant. Within this standard framework we have a loglikelihood function of the form [43]

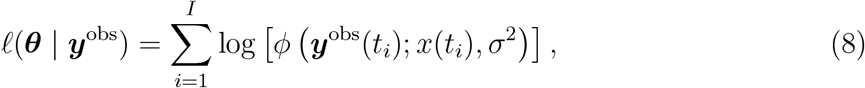

where *ϕ*(*x*; *µ, σ*^2^) denotes a Gaussian probability density function with mean *µ* and variance *σ*^2^. This commonly-used framework gives rise to the possibility of dealing with non-physical negative densities, especially in the situation where *p* is sufficiently small and either *k*_1_ or *k*_2_ is sufficiently large. Despite this potential pitfall, we note that working with additive Gaussian noise is very standard, especially in the systems biology literature [43], so we proceed with this standard approach for the moment, and in Sections 3.2 and 3.3 we provide concrete approaches for avoiding this complication.

With the loglikelihood function, Equation (8), numerical optimisation gives 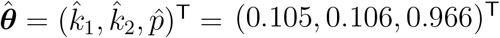, and the solution of the mathematical model evaluated with 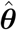 compares very well with the noisy data, as shown in Figure 1(a). To confirm that these parameter estimates are not identifiable we compute univariate profile likelihood functions for each parameter, as shown in Figure 1(b)–(d). For *k*_1_ and *k*_2_, as expected, the univariate profiles are perfectly flat, whereas the univariate profile for *p* is peaked about the MLE, and the approximate 95% confidence interval contains the true value. Since this model is non-identifiable we do not attempt to interpret biological mechanisms based upon our estimates of 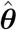 and nor do we attempt to make predictions from the non-identified model, instead we seek to simplify the model using a simple reparameterisation of the likelihood function. While it is possible to use the methods outlined in this work to make predictions from non-identifiable models [45], this involves working with user-defined bounds on the non-identifiable parameters rather than working with bounds defined by the curvature of the loglikelihood function.

**Figure 1:**
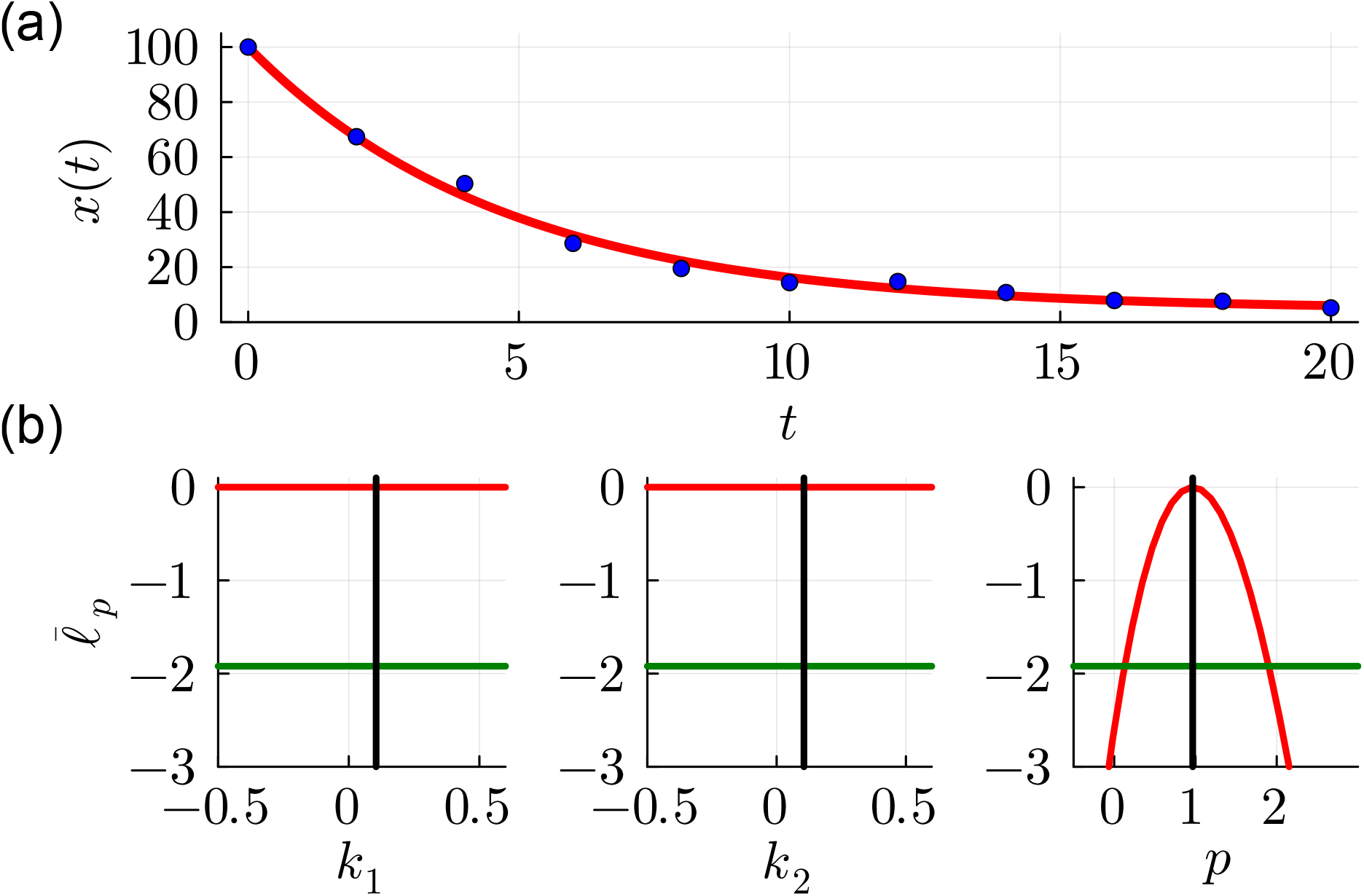
(a) Synthetic data (blue dots) obtained by evaluating the solution of Equation (6) at 11 equally-spaced values of *t* in the interval 0 ≤ *t* ≤ 20 and corrupting each value of *x*(*t*) with additive Gaussian noise with *σ* = 3. Synthetic data are generated with true parameter values ***θ*** = (*k*_1_, *k*_2_, *p*)^⊤^ = (0.1, 0.1, 1.0)^⊤^. The MLE solution (solid red) corresponds to 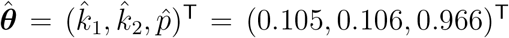. (b)–(d) Univariate profile likelihood functions for *k*_1_, *k*_2_ and *p*, respectively. Each profile shows 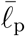 (red), together with the MLE (vertical black line) and the 95% threshold 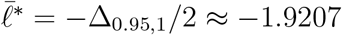 (horizontal green line).

Taking a standard approach to reparameterisation, we take the same data and mathematical model under the reparameterisation (*α*_1_, *α*_2_)^⊤^ = (*k*_1_ + *k*_2_, *p*)^⊤^ that is implied by the mathematical structure of the model or would be identified by standard structural identifiability software. The solution of the mathematical model evaluated at the MLE in Figure 2(a) compares well with the data. Importantly, the loglikelihood function in Figure 2(b) is characterised by a single, well-defined peak about the MLE, 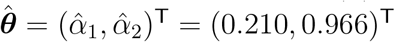. Both univariate profiles in Figure 2(c) contain a single well-defined peak about the MLE.

**Figure 2:**
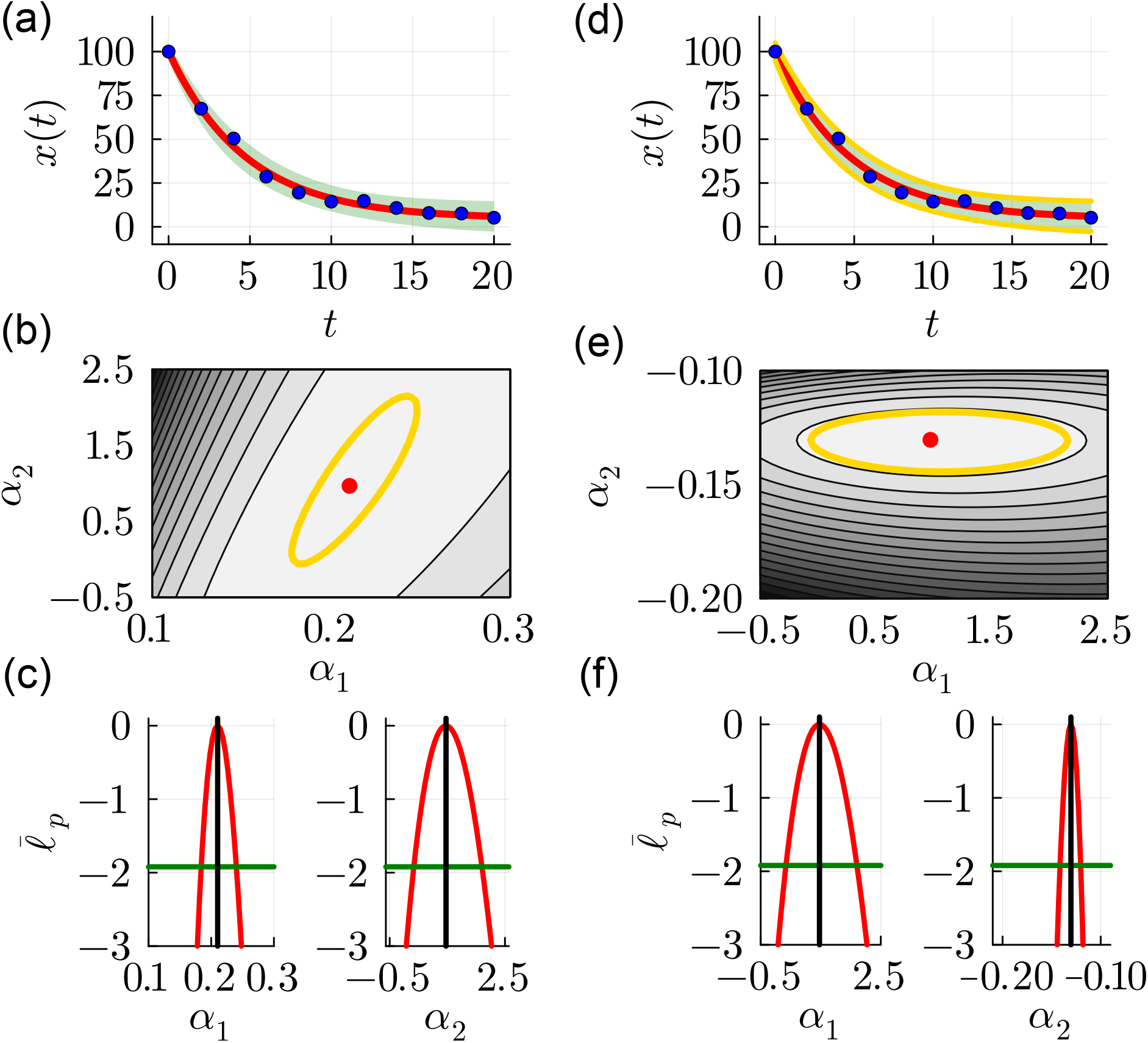
Results for the standard reparameterisation ***θ*** = (*α*_1_, *α*_2_)^⊤^ = (*k*_1_ + *k*_2_, *p*)^⊤^ in (a)–(c) are compared with results obtained using the eigendecomposition reparameterisation in (d)–(f). (a) Synthetic data (blue dots) from Figure 1 are superimposed with the MLE with 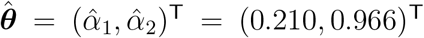 together with the 95% prediction interval (shaded green). (b) Normalised loglikelihood function superimposed with the MLE (red dot with 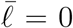). (c) Univariate profile likelihood functions for *α*_1_ and *α*_2_. (d) Synthetic data (blue dots) from Figure 1 are superimposed with the MLE with 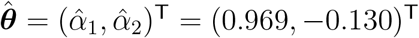 together with the exact (green shaded) and approximate 95% prediction intervals (solid gold curves). (e) Normalised loglikelihood superimposed with the MLE (red dots) (f) Univariate profile likelihood functions for *α*_1_ and *α*_2_ (red). (b) and (e) show the 95% threshold 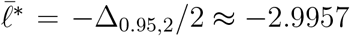 (gold curves). (c) and (f) show the MLE (vertical black line) and the 95% threshold 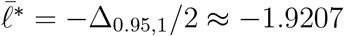 (horizontal green line).

For clarity we highlight the asymptotic 95% threshold, 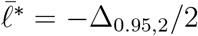 on the contour plot in Figure 2(b), indicating that estimates of *α*_1_ and *α*_2_ are positively correlated. This relationship between *α*_1_ and *α*_2_ makes intuitive mechanistic sense because it is possible to approximately match the same set of data by increasing (or decreasing) the total decay rate *α*_1_, while simultaneously increasing (or decreasing) the production rate *α*_2_.

To quantify and visualise how variability in the measurements translates into variability in estimates of ***θ***, and ultimately into variability in predictions, *x*(*t*), we use rejection sampling to obtain *M* = 1000 samples of ***θ*** within the region in Figure 2(b) for which 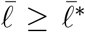 and construct the prediction interval shown in Figure 2(a). While the correlation structure inherent in this reparameterisation of the loglikelihood function makes intuitive sense, it means that sampling ***θ*** within the confidence set is complicated by this correlation structure.

An alternative approach is to consider a reparameterisation of the full ***θ*** = (*k*_1_, *k*_2_, *p*)^⊤^ parameter space by considering an eigendecomposition of the observed Fisher Information evaluated at the MLE. Here, with three parameters, the 3 × 3 observed Fisher Information has rank 2, with one zero eigenvalue. The two eigenvectors associated with the non-zero eigenvalues span the identifiable parameter space define a linear relationship between the eigenparameters and the model parameters, which for our data is

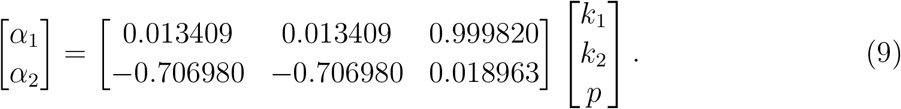

Here the elements of **v**_1_ are omitted since this eigenvector spans the non-identifiable parameter space. Equation (9) allows us to evaluate the eigenparameters (*α*_1_, *α*_2_)^⊤^ in terms of the original model parameters, (*k*_1_, *k*_2_, *p*)^⊤^, and we can re-write this relationship so that we can express the original parameters in terms of the eigenparameters. As we will demonstrate, working with the eigenparameters ***θ*** = (*α*_1_, *α*_2_)^⊤^ leads to a reduced and identifiable model that provides two advantages that become clear when we make predictions by sampling the loglikelihood function, as we will now explore.

The solution of the mathematical model evaluated at the MLE in Figure 2(d) compares well with the data. The loglikelihood function parameterised in terms of the eigenparameters, shown in Figure 2(e), is characterised by a single, well-defined peak about the MLE, 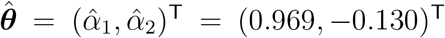. Both the corresponding univariate profiles in Figure 2(f) contain a single well-defined peak about the MLE. Estimates of 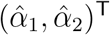 can be reinterpreted in terms of the model parameter *k*_1_, *k*_2_ and *p* using Equation (9).

A benefit of working in terms of the eigenparameters becomes clearer when we superimpose the 95% asymptotic threshold, 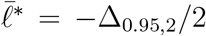 on the contour plot in Figure 2(e) which confirms that, as expected, there is lack of correlation between *α*_1_ and *α*_2_ in this orthogonal re-parameterisation. This lack of correlation means that mapping how the variability in ***θ*** impacts variability in predictions, *x*(*t*), is far simpler by rejection sampling. A second benefit is that the exact likelihood-based prediction interval (i.e. the green shaded region in Figure 2(d)) is accurately approximated by sampling from *K* = 2 univariate profile likelihood functions, and taking the union of the profile-wise prediction intervals to give the approximate prediction intervals (i.e. the region enclosed within the gold solid curves in Figure 2(d)). This accuracy of working with parameter-wise predictions is not always possible when the likelihood is correlated [37].

### 3.2 Morphogen gradient model

In this section we will work with a classical BVP that motivates us to move away from the standard additive Gaussian noise model that we used in Section 3.1 and dominates across the theoretical biology literature [43]. The motivation for avoiding additive Gaussian noise model is that we wish to preserve positivity of all data and predictions, and this is very important when working with data representing density and count data, for example when working with measurements of populations of cells or molecules. We proceed by considering a simple canonical reaction-diffusion equation that has often been used as a caricature model of the morphogen gradient formation during embryonic development that are thought to be associated with spatial patterning during morphogenesis [46, 47, 48]

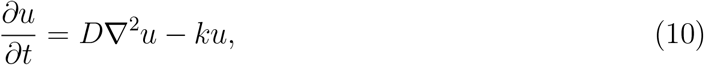

where *u* ≥ 0 represents a non-dimensional morphogen concentration. This model is often considered in a one-dimensional Cartesian geometry, where ∇^2^*u* simplifies to ∂^2^*u/*∂*x*^2^, and *u* = *u*(*x, t*). With the trivial initial condition *u*(*x*, 0) = 0 on *x* > 0, the dynamic morphogen gradient forms along the *x*-axis by applying a constant positive diffusive flux at the origin, *J* = −*D*∂*u/*∂*x* at *x* = 0. This simple model assumes that the morphogens undergo diffusion with constant diffusivity *D* > 0 as well as first-order linear decay with constant decay rate *k* > 0. To close the mathematical model, we assume the solution vanishes in the far field so that *u* → 0^+^ as *x* → ∞.

It is both mathematically convenient and biologically relevant to consider the long-time limit of the time-dependent model, 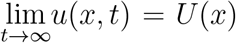, where *U*(*x*) is governed by the following BVP,

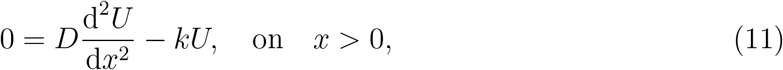

with d*U/*d*x* = −*J/D* at *x* = 0, and *U* → 0^+^ as *x* → ∞. At this point, even before solving the boundary value problem, we anticipate that this model will entail challenges of structural identifiability since the governing equation depends upon the ratio *D/k* and the boundary condition at *x* = 0 depends upon the ratio *J/D*. The solution of this BVP can be written as

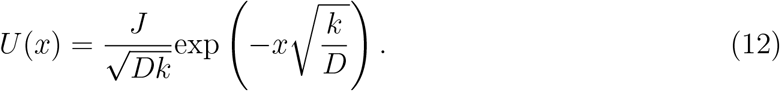

An important feature of this model is that we have *U* > 0 by definition, and the far-field boundary condition means that the density approaches zero for large *x*. This property means that the additive Gaussian noise model with constant variance is inappropriate as it would imply the possibility of negative densities, especially for large *x*. Another feature of this model that differs from the production-decay model is that the form of the solution in Equation (12) shows that the solution depends upon products and ratios of the three parameters, whereas the production-decay model involved parameter sums only. We deal with this by working with log(***θ***) = (log(*J*), log(*D*), log(*k*))^⊤^ which allows the eigendecomposition reparameterisation to be cast in terms of parameter products and ratios instead of sums and differences. This approach is a natural choice in many situations as key parameter combinations often take the form of monomials due to the Buckingham *π* theorem [49].

Data in Figure 3(a) are generated by evaluating the solution at 21 equally-spaced points on the truncated domain, 0 ≤ *x* ≤ 20, with true parameter values log(***θ***) = (log(*J*), log(*D*), log(*k*))^⊤^ = (log(1), log(1), log(0.1))^⊤^. To maintain *U* > 0, the solution at these points is corrupted with multiplicative log-normal noise with *σ* = 0.5 so that

**Figure 3:**
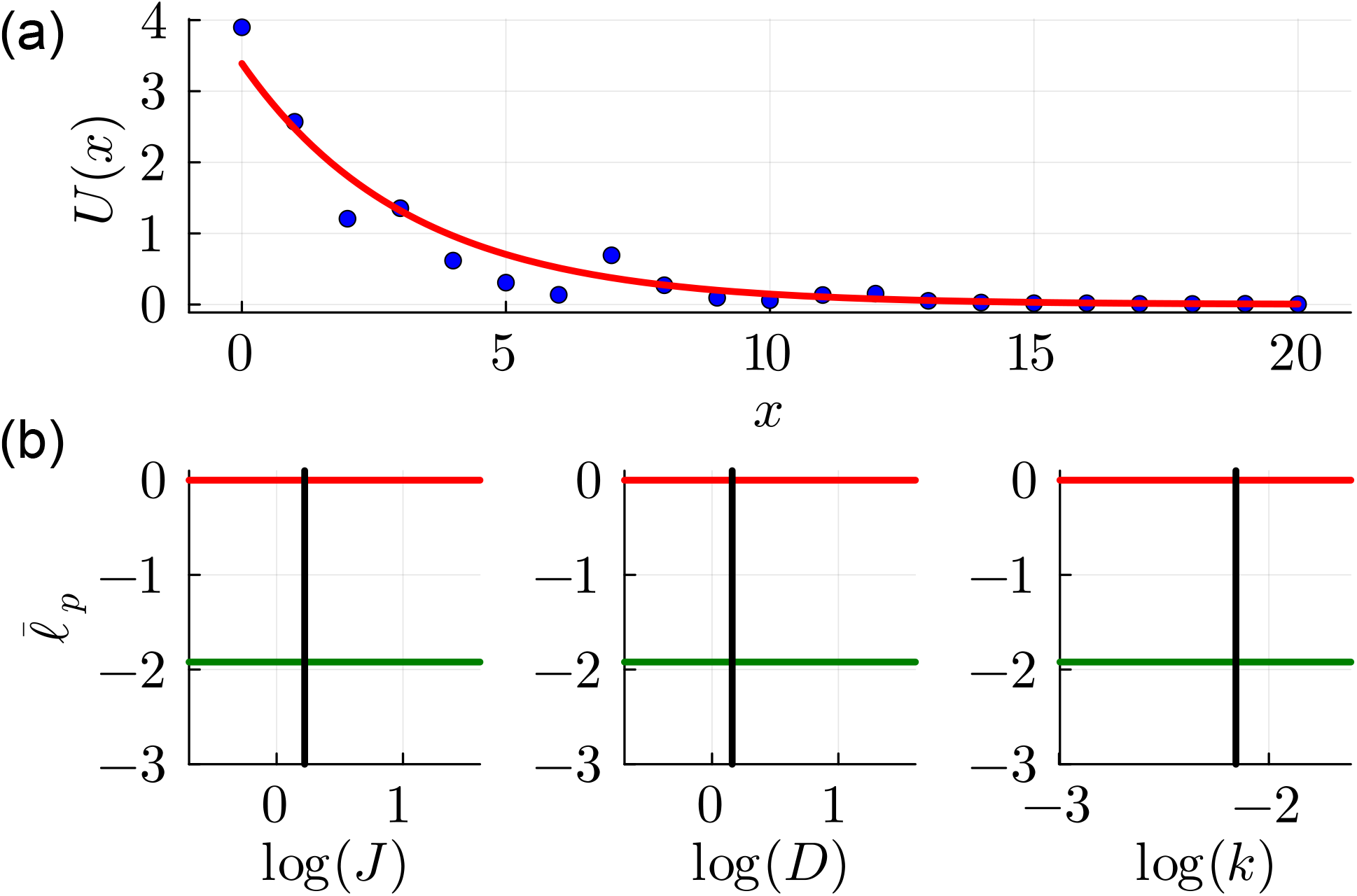
(a) Synthetic data (blue dots) obtained by evaluating the solution of Equation (11) at 21 equally-spaced values of *x* in the interval 0 ≤ *x* ≤ 20, and corrupting each value of *U*(*x*) with multiplicative log-normal noise with *σ* = 0.5. Synthetic data are generated with true parameter values ***θ*** = (*J, D, k*)^⊤^ = (1.0, 1.0, 0.1)^⊤^. The MLE solution (solid red) corresponds to 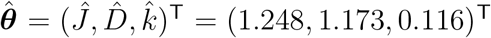. (b)–(d) Univariate profile likelihood functions for log(*J*), log(*D*) and log(*k*), respectively, showing that all three parameters are not identified by the data. Each profile shows 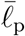 (red), together with the MLE (vertical black line) and the 95% threshold 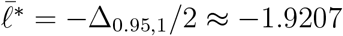 (horizontal green line).

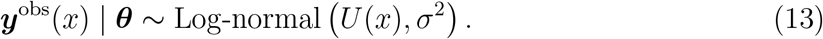

For this multiplicative noise model we see that the fluctuations in the data vanish for sufficiently large *x*, where *U* → 0^+^ in Figure 3(a). With this framework we have a loglikelihood function [43]

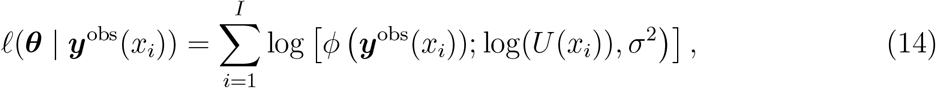

where *ϕ*(*x*; *µ, σ*) is the probability density function of the Log-normal(*µ, σ*^2^) distribution. Numerical optimisation gives 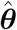, which we report here in terms of the original variables, 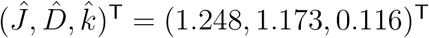. Superimposing *U*(*x*) evaluated at 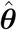 on the data indicates that the solution provides a good match to the data, but as with the production-decay model the MLE point estimate provides no insight into parameter identifiability. Parameter identifiability is assessed by constructing univariate profile likelihood functions for log(*J*), log(*D*) and log(*k*) in Figure 3(b). All three univariate profile likelihood functions are flat, confirming that the parameters are not well identified by this data. This is consistent with our previous observation that this model is structurally non-identifiable.

We will now illustrate how to undertake a model reduction using two approaches. First, taking a standard approach, we re-examine the same data and mathematical model under the reparameterisation that is motivated by examining the mathematical structure of the solution of the model, 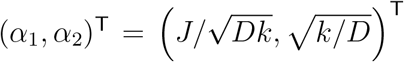. The solution of the mathematical model evaluated at the MLE in Figure 4(a) compares well with the data. The reparameterised loglikelihood function in Figure 4(b) is characterised by a single, well-defined peak at the MLE, 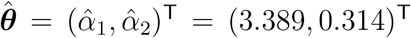, and both univariate profiles in Figure 4(c) each contain single well-defined peak at the MLE. As for the production degradation model, we highlight the 95% threshold, 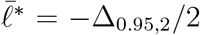 on the contour plot in Figure 4(b), which indicates that *α*_1_ and *α*_2_ are positively correlated. To understand how variability in the measurements propagates into uncertainty in our estimates of ***θ***, and ultimately into variability in predictions, *U*(*x*), we use rejection sampling to obtain *M* = 1000 samples of log(***θ***) within the closed contour in Figure 4(b) and construct the prediction interval shown in Figure 4(a). Here it is worth comparing the prediction intervals in Figure 2(c) and Figure 4(c). In the former, the additive Gaussian noise model with constant variance leads to a prediction interval whose width is approximately constant, whereas the width of the prediction interval for the log-normal noise model decreases with *U* so that the width of the prediction interval vanishes as *U* → 0^+^ in the far-field. This feature is important as it provides biologically meaningful predictions, whereas had we worked with the commonly-employed additive Gaussian model the predictions would have involved predicting negative densities [43], which is both biologically and physically unrealistic [44].

**Figure 4:**
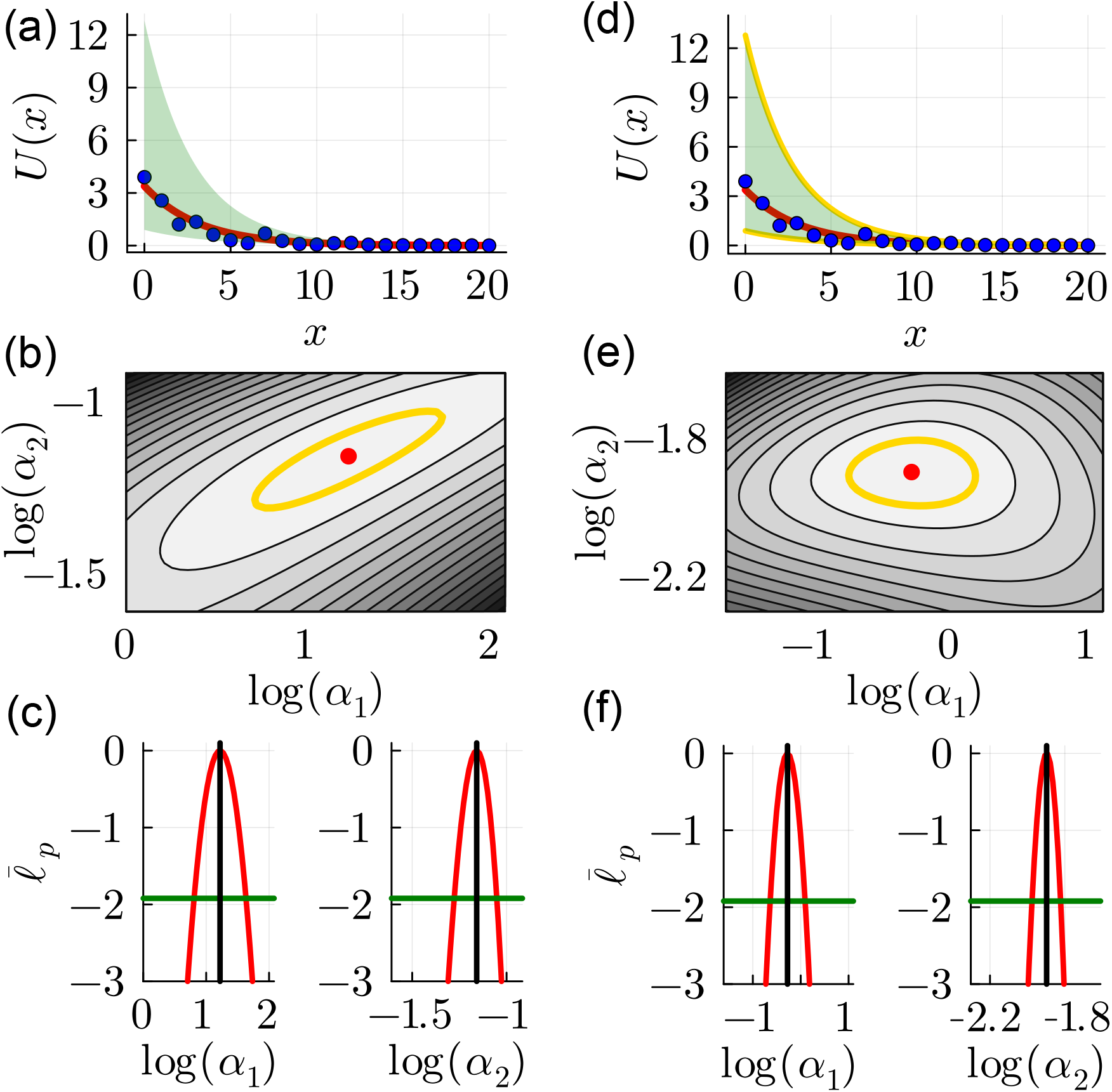
Results for the standard reparameterisation 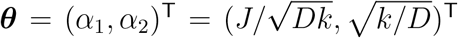 in (a)–(c) are compared with results for the eigendecomposition reparameterisation in (d)– (f). (a) Synthetic data (blue dots) from Figure 3 are superimposed with the MLE with ***θ*** = (*α*_1_, *α*_2_)^⊤^ = (3.389, 0.314)^⊤^ together with the 95% prediction interval (shaded green). (b) Normalised loglikelihood function with the MLE (red dot with 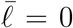). (c) Univariate profile likelihood functions for *α*_1_ and *α*_2_. (d) Synthetic data (blue dots) from Figure 3 are superimposed with the MLE with ***θ*** = (*α*_1_, *α*_2_)^⊤^ = (0.759, 0.150)^⊤^ together with the exact (green shaded) and approximate 95% prediction intervals (solid gold curves). (e) Normalised loglikelihood with the MLE (red dot) (f) Univariate profile likelihood functions for *α*_1_ and *α*_2_ (red). (b) and (e) show the 95% threshold 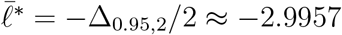 (gold curves). (c) and (f) show the MLE (vertical black line) and the 95% threshold 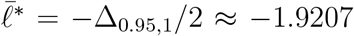 (horizontal green line).

We will now consider an eigendecomposition reparameterisation of log(***θ***) = (log(*J*), log(*D*), log(*k*))^⊤^ by computing the observed Fisher Information at the MLE. As with the production-decay model in Section 3.1, the 3 × 3 observed Fisher Information has rank 2 and one zero eigen-value. The components of the eigenvectors that span the identifiable parameter space defines a linear relationship between the logarithm of the eigenparameters and the logarithm of the model parameters,

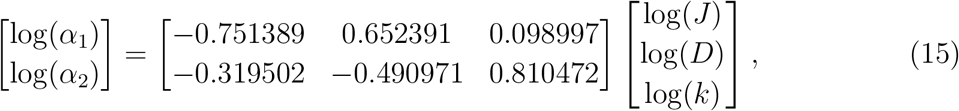

noting that elements of **v**_1_ are omitted since this eigenvector spans the non-identifiable parameter space. As before, working with the eigenparameters leads to a reduced and identifiable model that brings additional advantages in terms of making predictions by sampling the loglikelihood function, as we will now explore.

The solution of the mathematical model evaluated at the MLE in Figure 4(d) compares well with the data, and the loglikelihood function based on the eigendecomposition reparameterisation in Figure 4(e) is characterised by a single, well-defined peak at the MLE 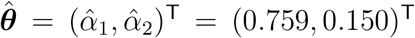. Both univariate profiles in Figure 4(f) contain a single well-defined peak at the MLE. As before, superimposing the 95% threshold, 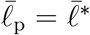 on the loglikelihood function in Figure 4(e) indicates that log(*α*_1_) and log(*α*_2_) are not correlated [50]. This lack of correlation structure again means that the computational task of sampling the loglikelihood function is simplified, and we find that the exact prediction interval obtained sampling the full loglikelihood function can be accurately approximated by sampling the two univariate profile likelihood functions and taking the union of the univariate predictions, as illustrated in Figure 4(d).

### 3.3 Cell invasion model incorporating cell cycle labelling

We now turn to a more practical problem of modelling spatiotemporal invasion of populations of cells that are often modelled using reaction-diffusion models in the context of a scratch assay, as illustrated in Figure 5(a)–(e). Scratch assays are performed by growing a monolayer of cells, creating a scratch in the monolayer, and then observing the population as individuals cells within the population undergo combined migration and proliferation, leading to a population-level closure of the scratched region [51, 52, 53]. These experiments are simple, fast, and inexpensive, which means they are routinely used as screening tools for exploring drug development or for understanding the population-level impact of genetic-level manipulations ahead of investing time and effort into more time-consuming, detailed and expensive experimental protocols. Since cell migration and cell proliferation are the dominant mechanisms, these experiments have often been modelled using the well-known Fisher-Kolmogorov model [54, 55], or generalisations of that model [56]. Formal structural identifiability analysis of PDE models is an area that is receiving considerable attention within the theoretical biology community [57, 58, 59]. In this case, however, the problem we consider relates to practical rather than structural identifiability [60].

**Figure 5:**
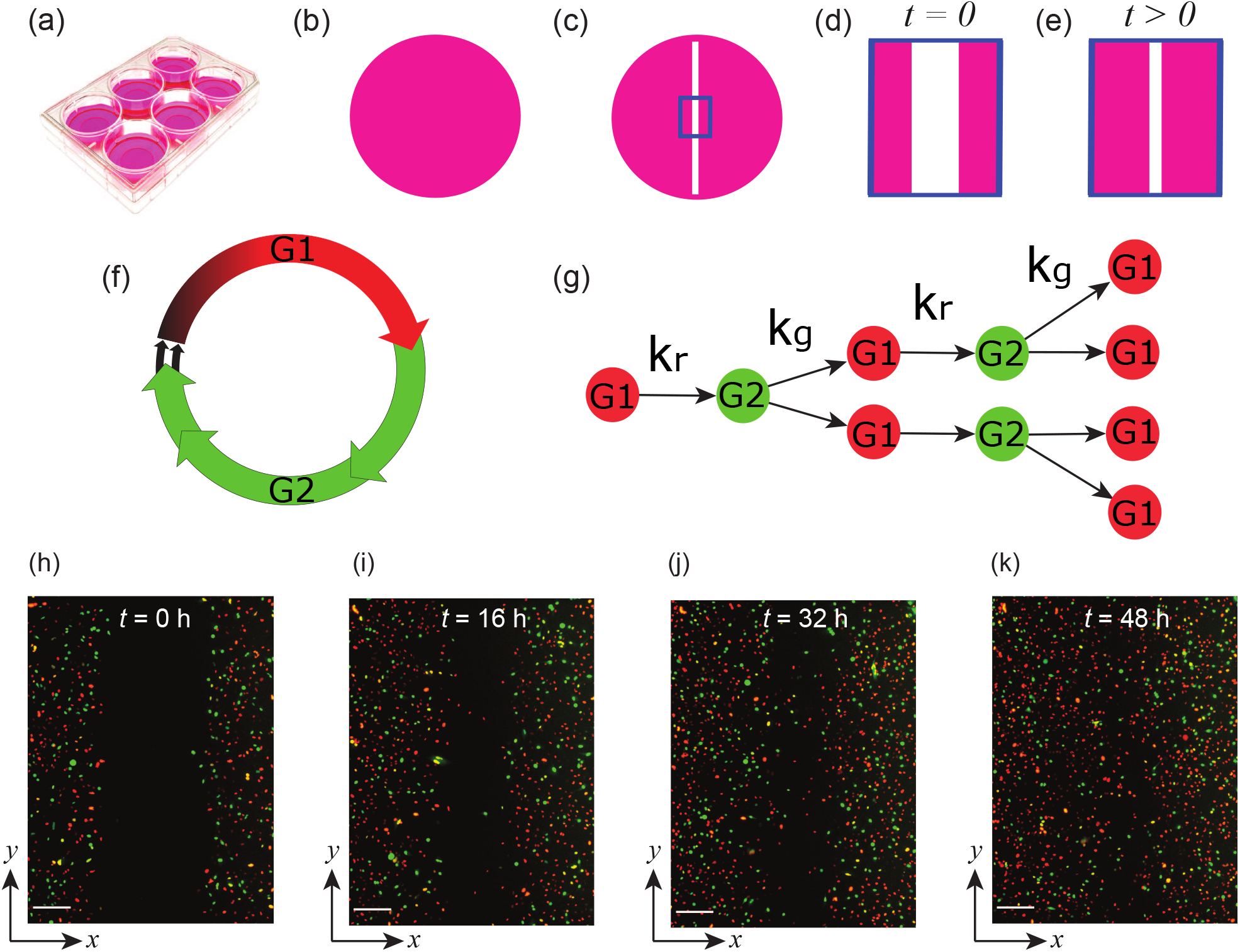
(a) Scratch assays performed in 6-well tissue culture places initiated with a low density, uniform monolayer of cells (pink shaded region). (b) A scratch into the monolayer is made (white region). (c) The field of view, 1296 *µ*m × 1745 *µ*m is observed between (d) *t* = 0 h (e) and *t* = 48 h. The cell cycle is labelled using fluorescent ubiquitination-based cell cycle indicator (FUCCI) [61, 62] so that cells in G1 phase fluoresce red and cells in S/G2/M phase fluoresce green (f). Freely-cycling cells in G1 phase (red) will transition into S/G2/M phase (green) at a rate *k*_*r*_ > 0. A freely-cycling cell in S/G2/M phase (green) will divide into two cells in G1 phase (red) at rate *k*_*g*_ > 0, provided there is sufficient space to accommodate the daughter cell (g). (h)–(k) Images of the scratch assay show the field of view at *t* = 0, 16, 32 and 48 h, where scale bars correspond to 200 *µ*m.

We will consider a set of scratch assay data reported by Vittadello et al. [63] performed with a human metastatic melanoma cell line, where individual cells contain a fluorescent marker so that cells in G1-phase of the cell cycle fluoresce red and individual cells within the S/G2/M phase of the cell cycle fluoresce green [61, 62]. For simplicity we will refer to those cells in the S/G2/M phase of the cell cycle as being in G2 phase. The cell cycle labelling technology enables the visualisation of both the spatial location and the cell cycle status of individual cells within an invading population of cells. A relatively simple PDE model, that can be interpreted as a generalisation of the Fisher-Kolmogorov model, was proposed by Vittadello et al. [63] to describe these experiments,

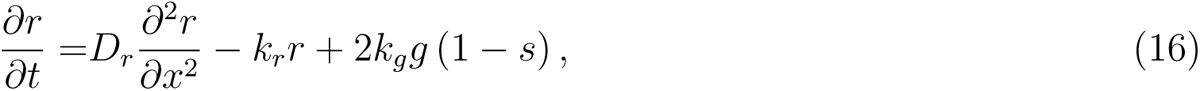

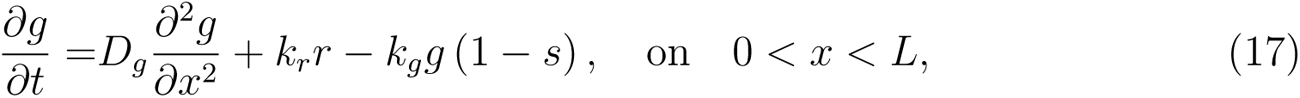

where *r*(*x, t*) ≥ 0 the non-dimensional density of cells in G1-phase that fluoresce red, *g*(*x, t*) ≥ 0 is the non-dimensional density of cells in G2-phase that fluoresce green, and *s*(*x, t*) = *r*(*x, t*) + *g*(*x, t*) is the total non-dimensional cell density. The parameters in this model are ***θ*** = (*D*_*r*_, *D*_*g*_, *k*_*r*_, *k*_*g*_)^⊤^, where *D*_*r*_ > 0 is the diffusivity of cells in G1-phase, *D*_*g*_ > 0 is the diffusivity of cells in G2-phase, *k*_*r*_ > 0 is the rate at which cells transition from G1-phase to G2-phase, and *k*_*g*_ > 0 is the rate at which cells transition from G2-phase to G1-phase. As illustrated in Figure 5(g), this second transition is associated with the production of two daughter cells, also in G1-phase, which is why the green–to–red transition is proportional to 2*k*_*g*_ in the source term in Equation (16). The factor of (1 − *s*) in this term is a parsimonious way to model the fact that the green–to–red transition is subject to contact inhibition since there needs to be sufficient space available to accommodate the new daughter cell produced by each individual division event. We note that the factor of (1 − *s*) is analogous to classical logistic growth model that also represents contact inhibition of proliferation by a quadratic nonlinearity [54, 55]. This one-dimensional mathematical model is appropriate to describe data from the experiments presented in Figure 5 because the geometry of the scratch within the uniform monolayer ensures that the macroscopic density varies only with horizontal position *x*, and time *t*. In contrast, the macroscopic density of both the red and green cell subpopulations is independent of vertical position [63].

Unlike the models in Sections 3.1–3.2 that could be solved analytically, Equations (16)–(17) must be solved numerically. In this work we take a method-of-lines approach by uniformly discretising the spatial domain, 0 *< x < L*, with constant mesh spacing *δx*, and then approximating the spatial derivatives in Equations (16)–(17) with a standard central difference approximation. The resulting system of coupled ODEs can then be solved numerically.

Data for this experiment are obtained by taking images, where each image has a width 1296 *µ*m and height 1745 *µ*m, and discretizing each image into a series of uniformly–spaced vertical strips. In our case we work with 24 strips, each of width 54 *µ*m. Counts of red and green cells within each strip, *R*^obs^(*x, t*) and *G*^obs^(*x, t*), respectively, are recorded for images at *t* = 0, 16, 32, 48 h. Assuming that these counts are associated with a horizontal position corresponding to the midpoint of each strip, our counts correspond to positions *x* = 0, 54, 104, … 1188, 1242 *µ*m. All counts can be converted into estimates of non-dimensional densities by dividing the raw counts by the strip area, and then normalising by a maximum packing density. Taking data at *t* = 0, linear interpolation provides an estimate of *r*(*x*, 0) and *g*(*x*, 0), which we use as initial conditions to solve Equations (16)–(17). Since the scratch is made in a uniform monolayer of cells, we apply zero net flux boundary conditions at *x* = 0 and *x* = *L* = 1242 *µ*m [63], which is consistent with the experimental protocol where the monolayer is uniform away from the location of the scratch. With *δx* = 5 *µ*m, this means that our numerical simulation of Equations (16)–(17) involves solving a system of 249 coupled nonlinear ODEs using DifferentialEquations.jl in Julia [64].

The count data for this experiment, shown in Figure 6, consists of non-negative integers that are bounded above by the maximum packing number of cells per strip, 𝒩. It is worth noting that this count data shows that cell counts per column are well below the maximum packing density. This is true even at the last time point in the experiments where the numbers of cells per column have more than doubled during the 48 hour duration of the experiment. Working with low density monolayers of cells is an intentional feature of these experiments because a key aim is to observe the individual location of cells and their cell cycle status as the experiment proceeds and the scratched region is recolonised [63]. Working with low density experiments is desirable because it avoids contact inhibition of proliferation that occurs under high density environments. Quantitatively, we have *s <* 0.2 for all *x* and *t* during the experiment, which means that we could model the same experimental data with a linear model by setting the (1−*s*) factors to unity in Equations (16)–(17). We do not follow this approach here as this simplified linear model leads to unrealistic infinite cell densities over longer timescales beyond the 48 hour duration of the current experiment.

**Figure 6:**
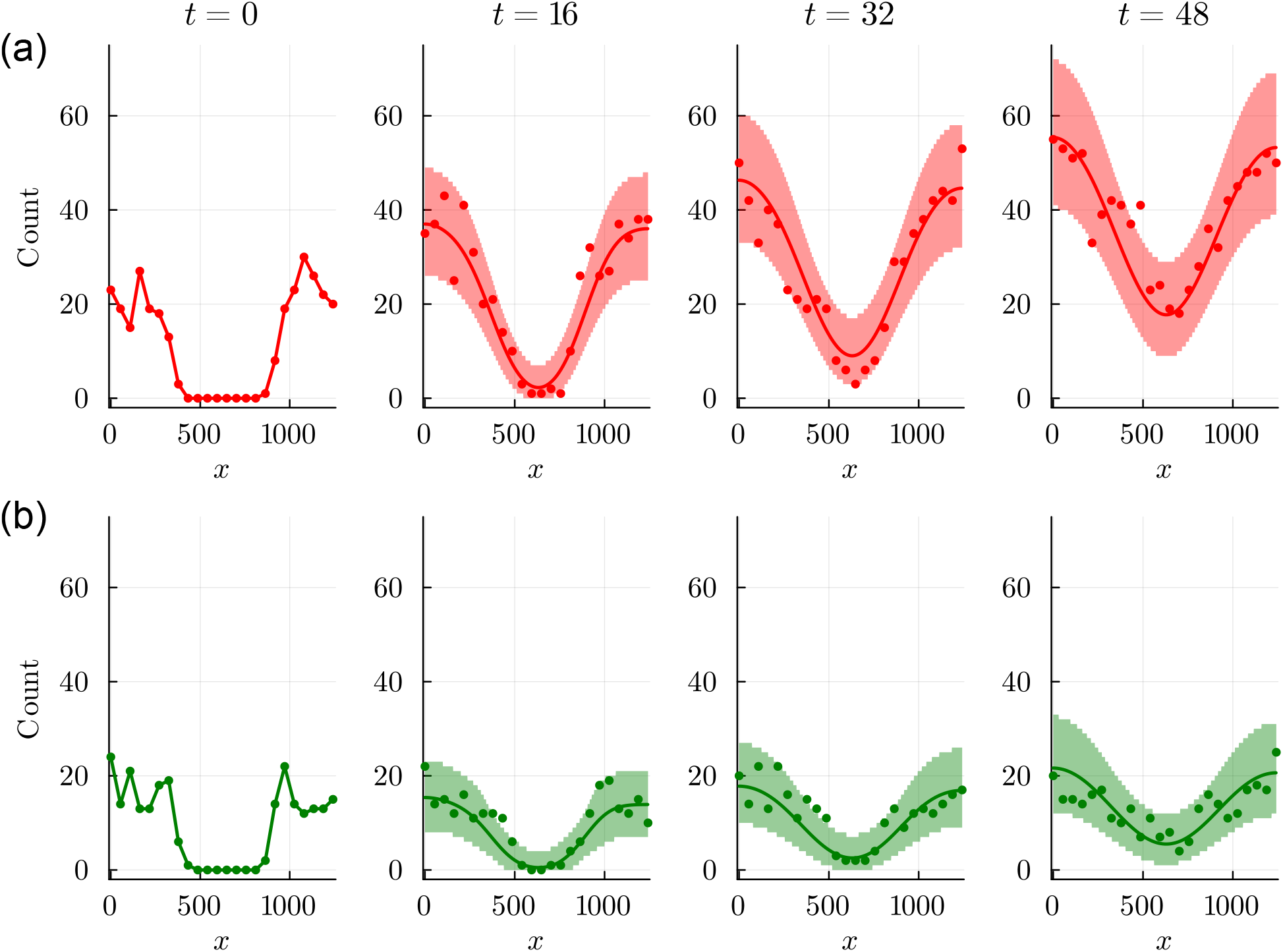
Scratch assay count data show the number of cells in each 54 *µ*m wide strip across the experimental images in Figure 1: (a) G1 phase (red dots) and (b) G2 phase (green dots) at *t* = 0, 16, 32, 48 hours, as indicated. Each plot shows the MLE solution of Equations (16)–(17) with 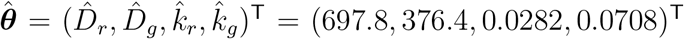. Each plot also includes a shaded prediction interval, red in (a) and green in (b), that are formed by taking *M* = 1000 random samples of ***θ*** and solving Equations (16)–(17) for each ***θ*** for which 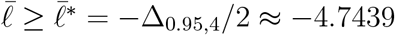 and then compute the prediction interval using 5% and 95% quantiles of the marginal distributions associated with multinomial noise model. All values of diffusivities have units of *µ*m^2^/h and all rates have units of /h.

For our initial condition, where *s*(*x*, 0) = *r*(*x*, 0) + *g*(*x*, 0) *<* 1 at all locations 0 *< x < L*, the numerical solution of Equations (16)–(17) leads to estimates of *r*(*x, t*) ∈ [0, 1] and *g*(*x, t*) ∈ [0, 1]. Within this setting we can interpret *r*(*x, t*) and *g*(*x, t*) as the fractions of the strip, centred at location *x*, that is occupied by cells in G1-phase and G2-phase, respectively. The finite size of cells, with approximate cell diameter 16 *µ*m, indicates that the maximum number of cells that can be placed into each strip is approximately 𝒩 = 377 [63]. This means in each strip we have *S*^obs^(*x, t*) = *R*^obs^(*x, t*) + *G*^obs^(*x, t*) total cells, and *E*^obs^(*x, t*) = 𝒩 − *S*^obs^(*x, t*) empty spaces that could accommodate cells of either type. To proceed, we interpret *R*^obs^(*x, t*) and *G*^obs^(*x, t*) as finite noisy samples from an underlying stochastic processes (i.e. the scratch assay experiment). These samples can be thought of as an approximation of some expected, noise-free measure of agent occupancy that is given by the solution of Equations (16)–(17). These measurements imply a multinomial loglikelihood function

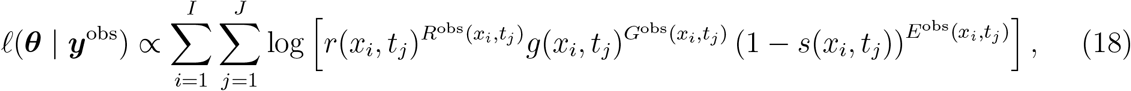

where the indices *i* and *j* refers to the locations in space and time, respectively, where count data are recorded [44]. This means that measurements are made at *I* = 24 spatial locations and at *J* = 4 instances in time. Given that each measurement involves recording counts of two cell types, we have a total of 2 × 24 × 4 = 192 measurements. Therefore, in this example, the length of ***y***^obs^ is 192. In practice we evaluate the multinomial loglikelihood function by setting the proportionality constant to unity. Since the initial condition for the PDE model is linearly interpolated from measured data, this means that data at *t* = 0 does contribute to 𝓁(***θ*** | ***y***^○^) in Equation (18). This means that we evaluate the loglikelihood by summing over contributions at *t* = 16, 32 and 48 h. Previous parameter estimation and identifiability analyses of PDE models of cell invasion typically take an approach by assuming that the data and PDE solution are related via additive Gaussian noise model [65, 60], here we take a different approach for two reasons: (i) count data are more naturally handled using a multinomial noise model; and (ii) standard additive Gaussian noise model implies unbounded predictions (e.g. negative counts), whereas the multinomial noise model leads to biologically-realistic non-negative discrete predictions that are bounded above by a carrying capacity density or maximum packing number [44].

To proceed we work with log(***θ***) = (log(*D*_*r*_), log(*D*_*g*_), log(*k*_*r*_), log(*k*_*g*_))^⊤^, and numerical optimisation gives, 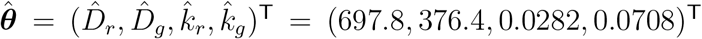. Throughout this document, all values of diffusivities are reported in units of *µ*m^2^/h and all rates are reported in units of /h. Plotting the solution of Equations (16)–(17) evaluated at 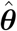, in terms of cell counts 𝒩*r*(*x, t*) and 𝒩*g*(*x, t*), shows that the solution of the mathematical model captures the broad trends in the experimental data, and provides a reasonable match to the noisy count data for both subpopulations across the 48-hour duration of the experiment.

To understand how well the data identifies the four parameters we construct four univariate profiles (see Appendix), and six bivariate profiles shown in Figure 7. The bivariate profiles indicate that five of six parameter pairs are reasonably well-identified by the data with the exception of the log(*ψ*) = (log(*D*_*r*_), log(*D*_*g*_))^⊤^ parameter pair. The other five bivariate profile likelihood functions are characterised by a single peak at the MLE, and the 95% confidence region relatively confined around the MLE without significant correlation. In contrast the log(*ψ*) = (log(*D*_*r*_), log(*D*_*g*_))^⊤^ bivariate profile likelihood function has a pronounced banana shape, indicating correlation between *D*_*r*_ and *D*_*g*_. This makes physical sense since the solution of the mathematical model with reduced (or increased) *D*_*r*_ can match the data reasonably well when *D*_*g*_ is increased (or reduced) to compensate. This is an example of a poorly-identified model where our data provides reasonable estimates of some model parameters (e.g. *k*_*r*_ and *k*_*g*_) whereas the others are poorly identified (e.g. *D*_*r*_ and especially *D*_*g*_). It is interesting to note that our data implies that our estimate of *D*_*r*_ is more precise than our estimate of *D*_*g*_. Data in Figure 6 confirms that these experiments involve more red than green cells, which is consistent with our observation that the diffusivity of the larger population, *D*_*r*_, can be estimated more precisely than the diffusivity of the smaller population, *D*_*g*_.

**Figure 7:**
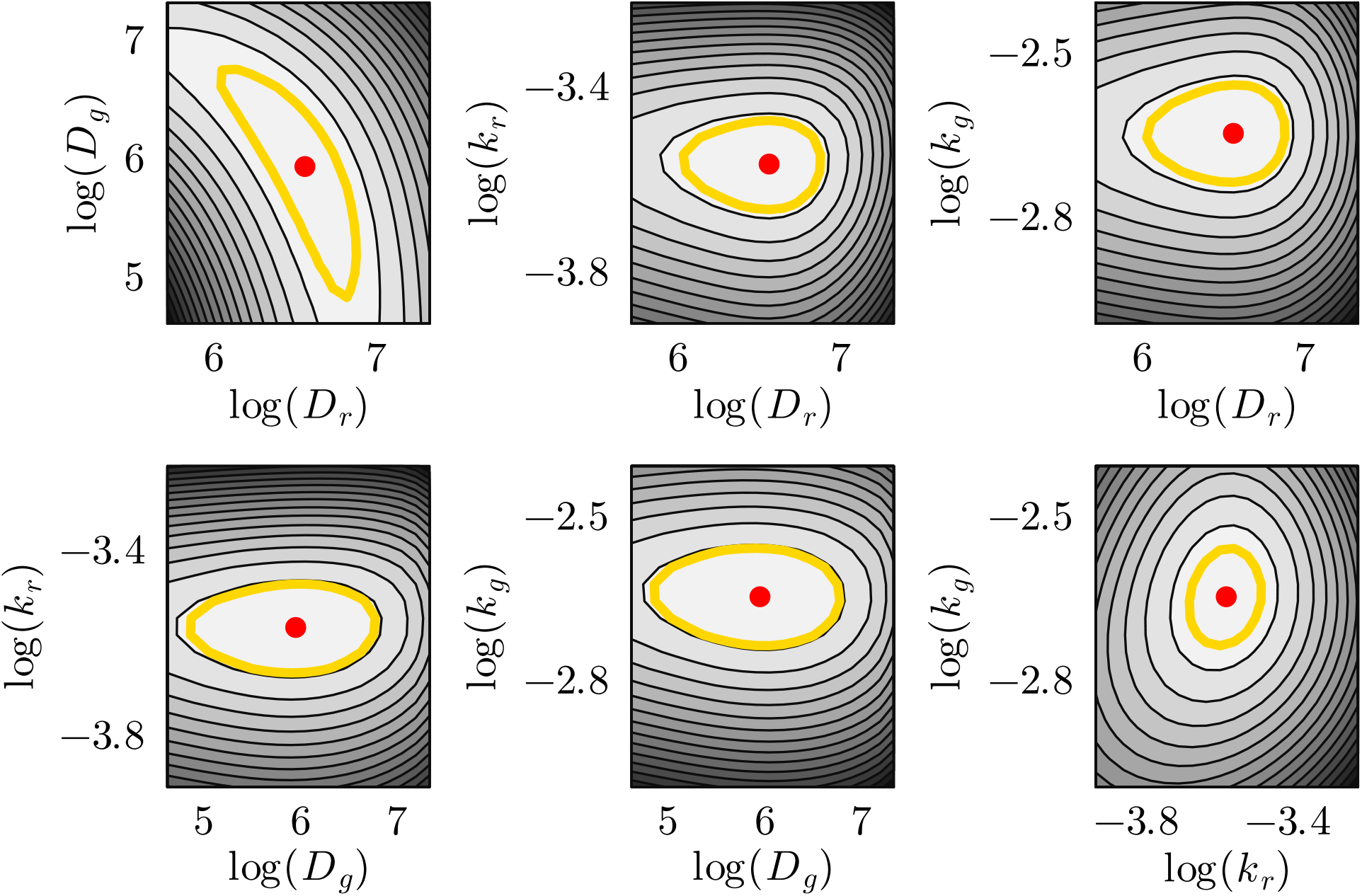
Six bivariate profile likelihood functions for the PDE model with log(***θ***) = (log(*D*_*r*_), log(*D*_*g*_), log(*k*_*r*_), log(*k*_*g*_))^⊤^. Each subplot shows one of six bivariate profile likelihood function 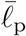, superimposed with the MLE, 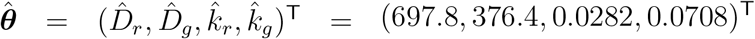 (red dot with 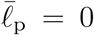). Each bivariate profile likelihood is superimposed with a curve illustrating the 95% threshold 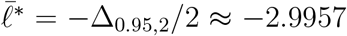 (gold solid curve). The greyscale differs in each subfigure with white corresponding to 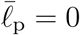, and darker shades correspond to decreasing 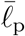. All values of diffusivities have units of *µ*m^2^/h and all rates have units of /h.

Regardless of the poor identifiability, we now explore the possibility of model predictions by using rejection sampling to obtain *M* = 1000 samples of log(***θ***) within in the region where 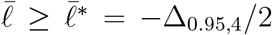. With these samples we construct the exact prediction intervals in Figure 6, where here we see the impact of the multinomial noise model since our predictions of the count data take the form of a series of staircase plots which reflect the fact that the multinomial noise model leads to non-negative integer predictions.

To address the issue or poor parameter identifiability we implement an eigendecomposition re-parameterisation, with the explicit aim of addressing the poor identifiability of *D*_*r*_ and *D*_*g*_. Evaluating the observed Fisher Information at the MLE shows that the 4 × 4 observed Fisher Information has rank 4, implying that the model is structurally identifiable, and that it is possible to reparameterise the model in terms of four eigenparameters. Again, noting the original parameterisation results in Figure 7 indicate that five of the six bivariate profiles contain a single peak at the MLE, and the 95% confidence regions are relatively constrained about the MLE, with the exception of the banana-shaped bivariate profile for log(*ψ*) = (log(*D*_*r*_), log(*D*_*g*_))^⊤^. This suggests that focusing on the two diffusivity parameters alone is a reasonable strategy since *k*_*r*_ and *k*_*g*_ are well-identified by the data. To proceed, we recall that the ordering of the non-zero eigenvalues gives the ordering of the four eigenparameters in terms of their identifiability. Therefore, we focus on the first eigenparameter, *α*_1_, which is the most identifiable of the four eigenparameters, and in this case can be written as

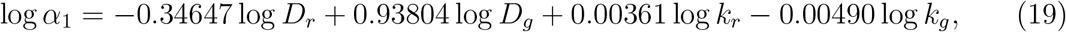

where we are working with a log-parameterisation with the aim that the reduced model can be written in terms of products or rations of the original model parameters. The form of the eigenparameter given by Equation (19) illustrates that *α*_1_ is relatively weakly-dependent on *k*_*r*_ and *k*_*g*_. This suggests an approximate reparameterisation, log *α*_1_ = −0.34647 log *D*_*r*_ + 0.93804 log *D*_*g*_, or 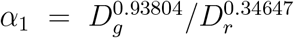, leading to an approximate reparameterisation log(***θ***) = (log(*α*_1_), log(*k*_*r*_), log(*k*_*g*_))^⊤^. Working with this reparameterisation, numerical optimisation gives 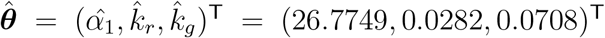. Plotting the solution of Equations (16)–(17) evaluated at 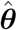 in terms of cell counts, 𝒩*r*(*x, t*) and 𝒩*g*(*x, t*), shows that the solution of the reduced mathematical model provides a good match to the noisy count data. To understand how well the data identifies the three parameters we construct three univariate profiles (see Appendix) and three bivariate profiles in Figure 8. The bivariate profiles show that each pair of parameters are well-identified by the data, and all three bivariate profiles display little correlation.

**Figure 8:**
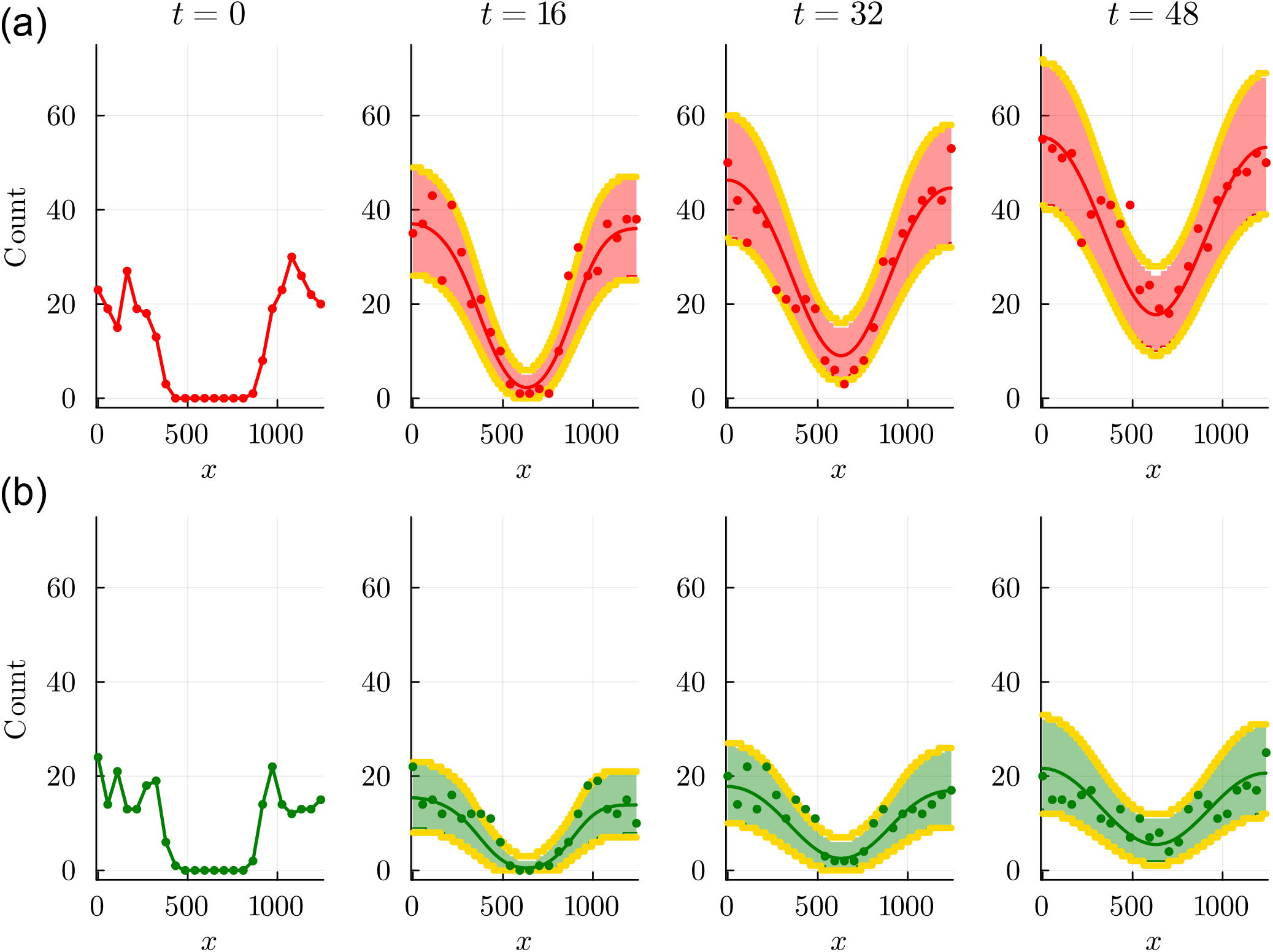
Scratch assay count data show the number of cells in each 50 *µ*m wide strip across the experimental images in Figure 6: (a) G1 phase (red dots) and (b) G2 phase (green dots) at *t* = 0, 16, 32, 48 hours, as indicated. Each plot shows the MLE solution of Equations (16)–(17) with 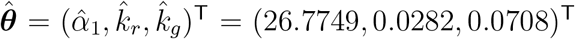. Each plot includes two different intervals. Shaded exact prediction intervals, red in (a) and green in (b), that are formed by working with *M* = 1000 samples from the full likelihood, whereas approximate prediction intervals formed taking *M* = 50 samples from each univariate profile likelihood and then taking the union across the three univariate profile likelihood functions are also superimposed (solid yellow curves). All values of diffusivities have units of *µ*m^2^/h and all rates have units of /h.

As with the previous eigendecomposition reparameterisations in Sections 3.1–3.2, the exact prediction interval obtained by densely sampling the full four-dimensional loglikelihood function is very accurately approximated by sparsely sampling the three univariate profile likelihood functions and taking the union of the univariate predictions, as illustrated in Figure 8.

## 4 Conclusion and future work

This article contains a series of implementation-focused computational explorations regarding reparameterisation of both structurally and practically non-identifiable mathematical models. Having the ability to implement a computational workflow to address the reparameterisation of non-identifiable models is important in the field theoretical biology where many standard mathematical modelling frameworks suffer from different forms of non-identifiability. Previous applications of model reparameterisation have focused on specific methods in terms of structural non-identifiability (e.g. algebraic and symbolic reparameterisation methods) separately from methods for practical non-identifiability (e.g. profile-likelihood-based approaches). Here we provide details of an easy-to-implement computational workflow that can be applied regardless of the source or form of non-identifiability. In terms of dealing with practical non-identifiability, previous applications of re-parameterisation via eigende-composition have focused on reduced models involving eigenparameters that take the form of products and ratios of original model parameters. Here, we extend this approach to deal with structurally non-identifiable models, and we illustrate how to obtain reduced models in terms of identifying eigenparameters in the form of products and ratios, or in the form of sums and differences of original model parameters. Working with eigenparameters in the form of products/ratios or sums/differences means that these eigenparameters are less interpretable than original model parameters, however the advantage of taking this approach is that the eigenparameters are well-identified by the available data, whereas the original parameters may not be.

To ensure that the approach is presented in a practical format we have demonstrated how to implement the method on a range of simple mathematical models that are very familiar across the field of theoretical biology. In particular, we apply the approach to: (i) a simple linear ODE model in the form of an IVP for a simple population dynamics problem; (ii) a second simple linear ODE model in the form of a BVP describing a classical morphogen gradient problem; and, (iii) a nonlinear coupled PDE model motivated by modelling spatiotemporal cell invasion problems. Across these three types of process models, we work with different kinds of noise models, namely: (i) additive Gaussian noise; (ii) multiplicative log-normal noise; and, (iii) a multinomial noise model suited to working with count data, respectively. Comparing the computational details of these three approaches confirms that the same framework can be applied to all three combinations of process models and noise models, thereby highlighting the versatility of the approach. This survey of different types of process models with different types of noise models avoids the pitfalls of focusing on a single type of mathematical model only (e.g. focusing only on ODE-based models with additive Gaussian noise). To ensure complete reproducibility of these approaches we provide simple user-friendly software, written in the open access Julia language so that all calculations can be repeated, and extended to other situations.

In each example, we focus on illustrating how the reparameterised model can be used to study model prediction, thereby allowing us to systematically examine how variability in data and observations propagates into variability in parameter estimates, and ultimately further propagating into variability in predictions and observations. This cascade of uncertainty propagation is important when theoreticians collaborate with experimental scientists as it provides us with an avenue for quantitatively understanding the how difference experimental designs and data collection protocols impact our ability to make projections and predictions via model calibration. In each case considered, we illustrate how to compute a prediction interval using the full likelihood function, noting that this can involve computational challenges owing to the potentially complicated shape of the likelihood function and the associated correlation structure. As we illustrate, a useful approximation is to work with a series of univariate profile likelihood functions and compute the prediction intervals in terms of the union of parameter-wise predictions. Working with parameter-wise prediction intervals brings the advantage of allowing us to explore the impact of each parameters, one-at-a-time, on the overall prediction interval. Since the reparameterised models developed using the eigenparameter approach leads to very simple uncorrelated likelihood functions, the union of the parameter-wise prediction intervals can be very accurate, and obtained using less computational effort than dealing with the full log-likelihood function.

While we have chosen to explore model reduction for a relatively broad class of mathematical models and noise models, we deliberately focused on working with continuum mathematical models written in terms of differential equations. Although we have not explicitly dealt with stochastic models, we note that stochastic mathematical models can be dealt with within the same framework by invoking a standard mean-field approximation where the mean behaviour of the stochastic model is represented by a differential equation-based framework [66].

There are several avenues for further exploration within the framework for reparameterisation. In the first two examples, focusing on synthetic data, we generate noisy data using a specified noise model with a single parameter, *σ*. In the subsequent inference, identifiability and prediction steps we then treated *σ* as a known constant, which is a standard approach based on the assumption that the noise parameter can be pre-estimated from the data or from numerical experiments [5]. In reality, we may also wish to estimate parameters in the noise model simultaneously with the parameters in the process models. This can be achieved by including *σ* in the vector of unknown parameters [67]. Another avenue for extending this work is to note that we have reparameterised non- and poorly-identifiable mathematical models in terms of eigenparameters that either take the form of products (ratios) or sums (differences) of original model parameters. We have not dealt with model reparameterisations in terms of a combination of products (ratios) and sums (differences) of original model parameters simultaneously. Another point for further exploration is the broad and important question of model misspecification. In the synthetic data examples presented here we always work with a mathematical model that was used to generate the noisy dataset, whereas in the cell invasion example we work with a mathematical model that was already known to provide a reasonable match to the observed data. We acknowledge that it is possible to model the cell invasion data using a different PDE framework that incorporates different types of terms describing cell migration (e.g. nonlinear diffusion [65]) and/or cell proliferation (e.g. different sigmoid growth models [67]). The broad aim of this work is to use a minimal but realistic process model, and it is relevant to note that even the simple model we work with leads to challenges involving issues around practical parameter non-identifiability. Therefore, we anticipate that modelling the same data with any kind of extended or more complicated process model would exacerbate these identifiability challenges. Therefore, we focus on working with a relatively simple model only where possible.

## Data Accessibility

Julia implementations within Jupyter notebooks for all computations are available on GitHub https://github.com/ProfMJSimpson/ModelReduction.

## Acknowledgements

I thank the MATRIX mathematical research institute for hosting a one-week residential workshop entitled “Parameter identifiability in mathematical biology” (September 2024) where initial work on this project took place. I also thank Ruth Baker and Sean McElwain for helpful feedback on draft versions of thi

## Appendix Univariate profile likelihoods for the PDE cell invasion model

In the main document we provide insight into the identifiability of parameters for the PDE model through a series of bivariate profile loglikelihood functions for the standard and reparameterised model in Figure 7 and 9, respectively. Here, in Figure 10–11 we provide the corresponding univariate profile likelihood functions. These additional results provide essentially the same information conveyed by the bivariate profile likelihood functions except that they do not provide any information about the correlation structure of the loglikelihood, which is why we prioritised the presentation and discussion of the bivariate profile likelihoods in the main document.

**Figure 9:**
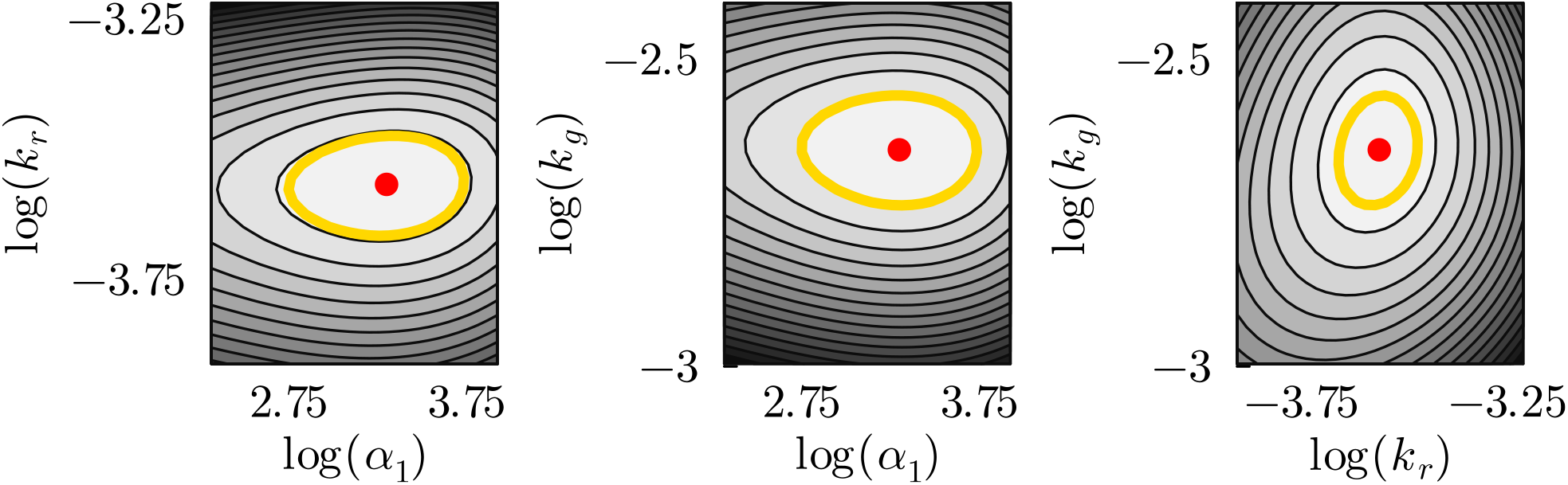
Three bivariate profile likelihood functions for the PDE model with the eigende-composition log parameterisation log(***θ***) = (log(*α*_1_), log(*k*_*r*_), log(*k*_*g*_))^⊤^. Each plot shows 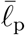 superimposed with the MLE, 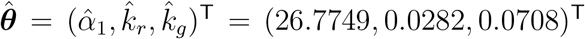 (red dot with 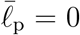). Each bivariate profile likelihood is superimposed with a curve illustrating the 95% threshold 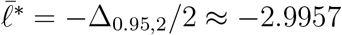 (gold solid curve). The greyscale differs in each subfigure with white corresponding to 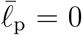 and darker shades correspond to decreasing 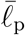. All values of diffusivities have units of *µ*m^2^/h and all rates have units of /h.

**Figure 10:**
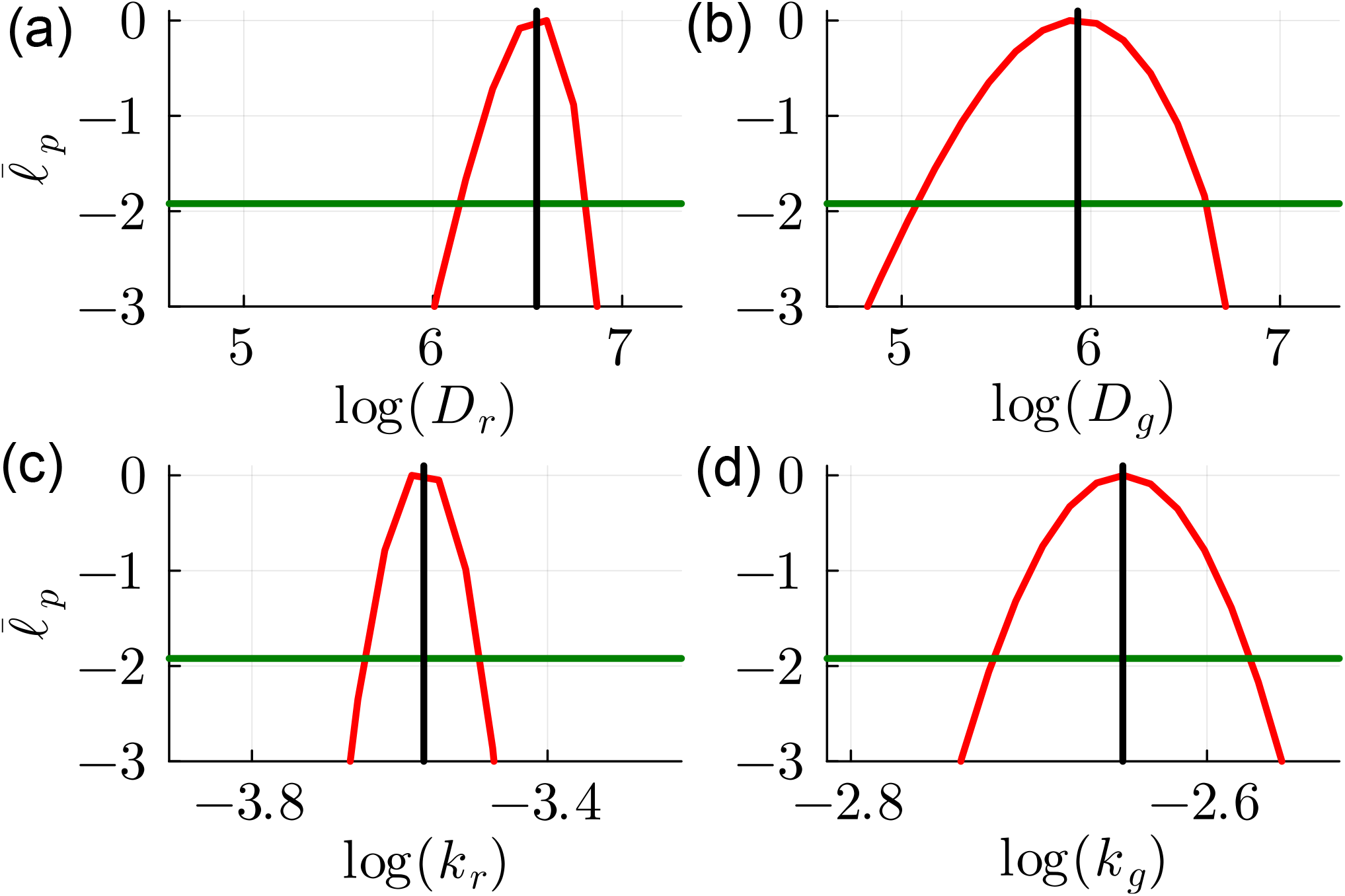
Four univariate profile likelihood functions for the PDE model with the log parameterisation log(***θ***) = (log(*D*_*r*_), log(*D*_*g*_), log(*k*_*r*_), log(*k*_*g*_))^⊤^. Each plot shows 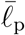 superimposed with the MLE (vertical black line), 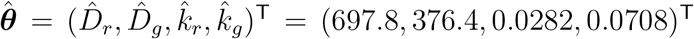 and the 95% threshold 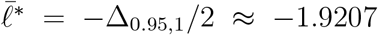 (horizontal green line). The MLE and 95% confidence intervals are: 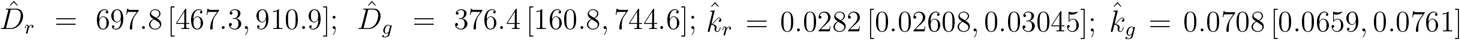. All values of diffusivities have units of *µ*m^2^/h and all rates have units of /h.

**Figure 11:**
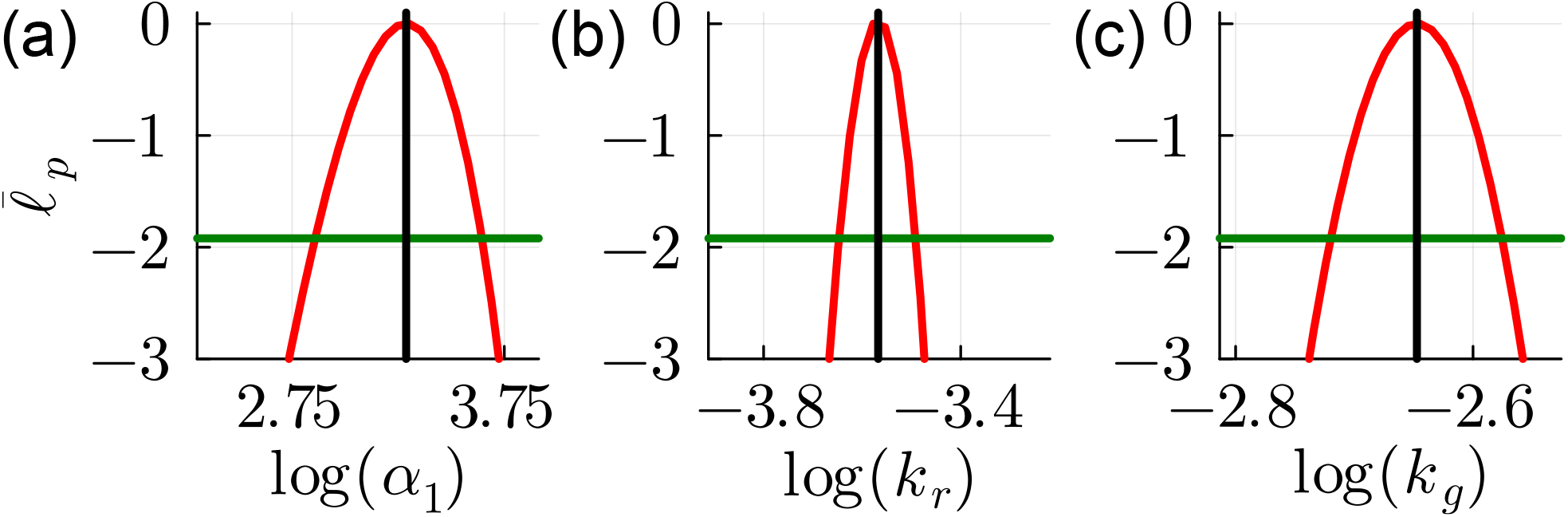
Three univariate profile likelihood functions for the PDE model with the eigen-parameter parameterisation log(***θ***) = (log(*α*_1_), log(*k*_*r*_), log(*k*_*g*_))^⊤^. Each plot shows 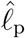 super-imposed with the MLE (vertical black line), 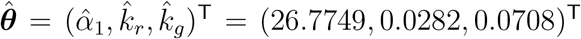 and the 95% threshold 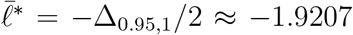 (horizontal green line). The MLE and 95% confidence intervals are: 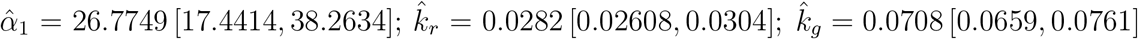. All values of diffusivities have units of *µ*m^2^/h and all rates have units of /h.

